# Potentiating TMEM16A channel function has no effect on airway goblet cells or bronchial and pulmonary vascular smooth muscle function

**DOI:** 10.1101/2020.04.17.046177

**Authors:** Henry Danahay, Roy Fox, Sarah Lilley, Holly Charlton, Kathryn Adley, Lee Christie, Ejaz Ansari, Camille Ehre, Alexis Flen, Michael J. Tuvim, Burton F. Dickey, Colin Williams, Sarah Beaudoin, Stephen P Collingwood, Martin Gosling

## Abstract

The calcium-activated chloride channel TMEM16A enables chloride secretion across several transporting epithelia, including in the airway where it represents a therapeutic target for the treatment of cystic fibrosis. Additional roles for TMEM16A have also been proposed, including enhancing goblet cell exocytosis, increasing goblet cell numbers and stimulating smooth muscle contraction. The aim of the present study was to test whether the pharmacological regulation of TMEM16A channel function, both potentiation and inhibition, could affect any of these proposed biological roles.

In vitro, a recently described potent and selective TMEM16A potentiator (ETX001) failed to stimulate mucin release from primary human bronchial epithelial (HBE) cells over a 24h exposure period using both biochemical and imaging endpoints. In addition, treatment of HBE cells with ETX001 or a potent and selective TMEM16A inhibitor (Ani9) for 4 days did not influence mucin release or goblet cell formation. In vivo, a TMEM16A potentiator was without effect on goblet cell emptying in an IL-13 driven goblet cell metaplasia model.

Using freshly isolated human bronchi and pulmonary arteries, neither ETX001 or Ani9 had any effect on the contractile or relaxant responses of the tissues. In vivo, ETX001 also failed to influence either lung or cardiovascular function when delivered directly into the airways of telemetered rats.

Together, these studies do not support a role for TMEM16A in the regulation of goblet cell numbers or mucin release, or on the regulation of airway or pulmonary artery smooth muscle contraction.

## Introduction

The TMEM16A protein (anoctamin-1) is now recognised as a calcium activated chloride channel (CaCC) and is the native CaCC in a variety of cell types including secretory epithelia, smooth muscle and neurones (Pedemonte & Galietta, 2014). The long-recognised functions of the CaCC in these cell types range from the regulation of ion transport and fluid homeostasis to the control of contractile responses in excitable tissues. With the identification of TMEM16A as the CaCC (Caputo et al., 2008; Schroeder et al., 2008; Yang et al., 2008), genetic models and pharmacological modulators have begun to introduce an expanding list of biological roles for this protein.

With respect to the function in the airways, Caputo and colleagues (2008) demonstrated that TMEM16A was the CaCC responsible for vectorial chloride transport across the bronchial epithelium and that both expression and function were upregulated by the Th2 cytokines IL-4 and IL-13. Enhancing TMEM16A-mediated chloride secretion thereby represents a therapeutic approach to hydrating the hyper-concentrated mucus in the airways of patients with a variety of diseases associated with mucus obstruction including cystic fibrosis (Kunzelmann et al., 2012; Mall & Galietta, 2015; Sondo et al., 2014). In support of this concept, a novel low molecular weight potentiator of TMEM16A, ETX001, has been demonstrated to promote airway mucosal hydration and to promote mucus clearance in vivo (Danahay et al., 2020).

Of further relevance to lung physiology, it has been proposed that TMEM16A can positively regulate both mucin secretion from goblet cells and the IL-13-dependent formation of goblet cells (Benedetto et al., 2019; Huang et al., 2012; Zhang et al., 2015; Lin et al., 2015; Qin et al., 2016) although a recent report challenges some of these observations (Simões et al., 2019). Additionally, there are several reports indicating that TMEM16A positively regulates the contractile responses of murine, guinea pig and human airway smooth muscle in vitro and murine airways in vivo (Danielsson et al., 2015, 2020; Huang et al., 2012; Miner et al., 2019). TMEM16A blockers, including niclosamide and benzbromarone, have been demonstrated to relax airway smooth muscle, whilst a reported TMEM16A activator, E_act_, induces contractile responses. To this end it has been suggested that blocking TMEM16A therapeutically may provide a greater overall benefit in the lungs of patients, by simultaneously bronchodilating the airways and reducing the burden of mucus, versus the benefit of enhancing TMEM16A function and improving mucosal hydration and mucus clearance (Kunzelmann et al., 2019). Similarly, enhancing the activity of TMEM16A with agents such as ETX001 could potentially increase mucus secretion and goblet cell numbers and cause bronchoconstriction which may represent safety issues in the clinic.

The aim of the present study was therefore to test the hypothesis that enhancing the activity of TMEM16A using potentiators of channel function, such as ETX001, would induce bronchoconstriction, airway mucus secretion, increase goblet cell numbers and alter pulmonary vascular tone. Using a variety of model systems, including primary human bronchial epithelial cells, in vivo airway mechanics and freshly isolated human bronchi and human pulmonary arteries, we demonstrate that ETX001 and related TMEM16A potentiators do not induce changes in airway smooth muscle function, do not influence either mucus secretion or the formation of goblet cells, and do not influence human pulmonary artery smooth muscle contraction. The observed lack of activity of the potent and selective TMEM16A inhibitor Ani9 (Seo et al., 2016) in these same airway models further challenges the conclusions of the published studies and a role for TMEM16A in these processes.

## Methods

Unless otherwise stated, all chemicals were purchased from Sigma (UK) and cell culture reagents from Life Technologies (UK).

### Cell culture

HBE were provided by Dr Scott Randell from both the University of North Carolina Chapel Hill collection and the Cystic Fibrosis Foundation Therapeutics repository. Cells were cultured at air-liquid interface as previously described (Fulcher et al., 2005; Coote et al., 2015). In brief, following an expansion step on plastic, HBE were seeded onto either Snapwell or Transwell inserts (Costar, UK) in submerged culture followed by 2 weeks at air-liquid interface in either defined ALI media (Fulcher et al., 2005) or DMEM:F12 media supplemented with Ultroser G (2% v/v; Pall, France) (Coote et al., 2015). At all stages of culture, cells were maintained at 37°C in 5% CO_2_ in an air incubator.

### Ion transport

HBE that had been cultured on Snapwell inserts for 2 weeks at air-liquid interface in Ultroser G-based media, were treated with IL-13 (Peprotech, UK) for 48-96h (10 ng/mL) prior to use in functional studies, to upregulate TMEM16A expression. In some studies, HBE were also treated with either ETX001 (1 μM) or vehicle (0.01% DMSO) for 96 h whilst in culture.

HBE were mounted in Ussing chambers in symmetrical Ringers solutions and voltage clamped to 0 mV as previously described (Coote et al., 2015). Following the addition of amiloride to inhibit the ENaC-mediated short-circuit current, the SERCA-pump inhibitor cyclopiazonic acid (CPA) or UTP were added to the cells to elevate [Ca^2+^]_i_ levels and to enable the efficacy of ETX001 to be evaluated. In the CPA-format, an EC_20_ concentration of the SERCA-pump inhibitor was added to the cells to induce a stable, TMEM16A-mediated current response, upon which a concentration-response to ETX001 could be generated, as previously described (Danahay et al., 2020). In the UTP format, increasing concentrations of UTP were added cumulatively and the area-under-the-curve was quantified to determine the magnitude of the anion secretory response.

### Mucus secretion

In *vitro:* CF-HBE that had been cultured in defined ALI media (Fulcher et al., 2005) were used for these studies. Cells from 2 CF-HBE donor codes, both homozygous for ^F508del^CFTR were used for these studies. Half of the inserts were treated with IL-13 (10 ng/mL) for 48h to increase the expression of TMEM16A and MUC5AC. After washing the apical surface to remove any accumulated mucus, cells were treated with vehicle (0.1% DMSO) or ETX001 (1 μM) for 24h. After treatment, cell washings (0.5 mM tris (2-carboxyethyl) phosphine in saline; 150 μL/insert; 30 min) were collected from ≥12 inserts per donor code. Phorbol 12-myristate 13-acetate (PMA; 300 nM) was used as a positive control to confirm the capacity to increase goblet cell exocytosis. Washings were separated by electrophoresis and probed with antibodies directed against MUC5AC (mouse monoclonal 45M1, Invitrogen) and MUC5B (rabbit polyclonal UNC414, provided by Ehre laboratory). Signal intensity was analysed using the LiCor Odyssey software and was normalised to an untreated control group. To assess any effects of treatment on mucin granule numbers in goblet cells, unwashed inserts (n=3) were fixed (osmium perfluorocarbon) and were processed for transmitted electron microscopy (TEM) as previously described (Ehre et al. 2012). Briefly, inserts were carefully fixed in 2.5% glutaraldehyde / 2% paraformaldehyde in 0.1M sodium cacodylate buffer at 37°C. Fixed samples were embedded, polymerized, serially sectioned via a TEM grid and ultrathin sections (~80 nm) were stained prior to imaging. The entire length of cell inserts was imaged using a JEOL 1230 transmission electron microscope at 8000x magnification. The average number of granules per field of view were counted across the entire length of each insert. A one-way ANOVA was used to test for differences between treatment groups.

In *vivo:* Airway goblet cell metaplasia was induced in C57bl6 mice by intrapharyngeal IL-13 instillation (4 × 2 μg in 40 μL dosed on days 1, 3, 5 & 13) as previously described (Zhu et al., 2015). On day 15, mice were administered either vehicle or a TMEM16A potentiator. To dose ETX001 directly into the airways requires the compound to be formulated as a suspension. To avoid any potential artefact induced by particulate instillation into the lungs in this model, a metabolically stable analogue of ETX001, ETX004, was dosed to mice by intraperitoneal injection. Doses of ETX004 were selected to achieve systemic exposures that were in excess of those required to fully engage the target (Figure 3A; Figure S1; Supplemental Methods for pharmacokinetic study details). At 2.5h after vehicle (5% NMP/95% HPCD [20%]) or compound dosing, mice were exposed to an aerosol of either normal saline or ATP (100 mM in saline) for 20 min. At 10 min after completion of the aerosol dosing, animals were euthanised and lungs fixed, processed and stained with PAFS as previously described (Evans et al., 2004). Images were acquired using an upright microscope (Olympus BX 60) with a ×40 NA 0.75 objective lens, and intracellular mucin was measured around all the circumferential section of the axial bronchus using ImagePro (Media Cybernetics) (Jaramillo et al., 2019). Data are presented as the epithelial mucin volume density, signifying the measured volume of mucin overlying a unit area of epithelial basal lamina, derived as described previously (Evans et al., 2004). Images were analysed by investigators blinded to treatment.

### Goblet cell formation

Following 14 days of differentiation at air-liquid interface in Ultroser-G based media, HBE were treated with either vehicle (0.1% DMSO), Ani9 (10 μM), ETX001 (1 μM) or the combination of ETX001 and Ani9. These study groups were replicated both in the presence and absence of IL-13 (10 ng/mL) treatment. All treatments were refreshed at 48 hours and maintained to 96 hours. At 96 hours, wells were fixed in 4% formaldehyde and were stained with antibodies to MUC5AC (45M1; Thermo Fisher, UK) and acetylated α-tubulin (6-11B-1; Sigma, UK) as previously described (Danahay et al., 2015) with secondary antibodies to enable fluorescence detection of both proteins (Alexa Fluor; Thermo Fisher, UK). The MUC5AC+ stained area was visualised using a Zeiss Axiovert epifluorescence microscope with a motorised stage that was used to image the same 9 regions of interest on each insert. Image J was used to quantify the MUC5AC+ and stained area per insert which was normalised to the vehicle control group. The process was repeated for the acetylated α-tubulin+ stained area. A one-way ANOVA with post-hoc Sidak test was used to enable statistically significant differences between specific groups to be identified.

### Wire myography

Non-transplantable fresh lungs from 3 organ donors were used for these studies. Donors with a history of asthma, COPD, emphysema, lung cancer, cystic fibrosis, pulmonary fibrosis, pulmonary hypertension or pneumonia were excluded. Airways and pulmonary arteries were dissected free from the surrounding parenchyma and used for wire myography studies as previously described (Miner et al., 2019). In brief, bronchi and arteries were cut into 2 mm rings and mounted under isometric conditions in 5 mL myographs containing a physiological salt solution, aerated with 95% O_2_ / 5% CO_2_ at 37°C. Airway preparations also had indomethacin (5 μM) included in the salt solution. Following stabilisation, tissue viability was assessed by challenging airways with carbachol (10 μM) and isoprenaline (10 μM) to confirm a bronchoconstrictor and dilation response respectively. Airways that failed to respond were excluded. The viability of human pulmonary arteries was similarly confirmed using a high K+ (62.5 mM) challenge to induce vasoconstriction. In addition, U44619 (100 nM) and acetylcholine (10 μM) were used to test endothelial integrity with constrictor and relaxation responses respectively. Following the viability assessment, all preparations were washed, and tension returned to starting levels. U46619 (100 nM) was added to pulmonary arteries selected for relaxation study. Test items were added to the preparations in increasing concentrations from 0.1 nM to 10 μM. At the completion of treatment, theophylline (2.2 mM) was added to the airway preparations to fully relax the tissue. A two-way ANOVA was used to test for significant differences between the test and vehicle treated groups.

### Lung and cardiovascular function in vivo

Male Sprague-Dawley rats (500-625 g) had previously been implanted with a HD-S10 telemetry transmitter (Data Sciences International, USA), with the pressure catheter positioned in the ascending aorta in order to measure blood pressure. A total of 10 animals were allocated to study, from which 8 were chosen, based on blood pressure signal and values. At weekly intervals, each animal received administration of either vehicle (0.05% Tween-80 in normal saline) or ETX001 (0.1 – 2.5 mg/kg) by intra-tracheal instillation whilst under isoflurane-induced anaesthesia.

Respiratory parameters were measured using whole body, bias flow plethysmography equipment designed by Buxco Research Systems (Data Sciences International, USA). Respiratory parameters (respiratory rate, tidal volume and minute volume) were derived from the changes in pressure associated with the warming and humidification of the air breathed in by the animal. This was monitored by specific probes located within the plethysmograph chambers. Bias flow (room air) was set at approximately 2.5 L/min. On one occasion prior to the first day of dosing, the animals were habituated to the plethysmographs for approximately 2h. On each test session, the animals were placed into plethysmograph chambers for pre-dose recording of respiratory parameters (respiratory rate, tidal volume, minute volume) for approximately 1h. Animals were then removed from the plethysmograph chambers and returned to their home cages. The animals were then dosed and returned to the plethysmograph chambers and continuously recorded for approximately 4h post-dose. Respiratory rate, tidal volume and minute volume were reported every 30 min up to 4h postdose. Time 0 was the mean value of the last 20 min of data recorded during the 1h pre-dose period. All post-dose time points were the mean of ten min recordings around each time point (e.g. 10 min before and 10 min after) with the exception of the 4h timepoint, which was the mean value of the last 20 min of data recorded.

Cardiovascular parameters (systolic, diastolic and mean blood pressure and heart rate derived from blood pressure) and core body temperature were recorded from the implanted transmitter using a dedicated Dataquest OpenART data capture system linked with a Ponemah data analysis system (Data Sciences International, USA).

For the respiratory and cardiovascular data (including core body temperature) each post-dose timepoint and parameter was analysed separately using analysis of covariance (ANCOVA). Factors in the model included day of measurement and treatment. Pre-dose data was included in the model as a covariate. ETX001 treatment was compared to vehicle treatment using Williams’ test.

### Study approval

The in-life experimental procedures undertaken in this study were subject to either the provisions of the United Kingdom Animals (Scientific Procedures) Act 1986 Amendment Regulations 2012 (the Act) or in accordance with the protocol approved by the Institutional Animal Care and Use Committees of The University of Texas MD Anderson Cancer Center (00001214-RN01). The number of animals used was the minimum that is consistent with scientific integrity and regulatory acceptability, consideration having been given to the welfare of individual animals in terms of the number and extent of procedures to be carried out on each animal.

## Results

### Results Epithelial ion transport

As previously described (Danahay et al., 2020), ETX001 (1 μM) potentiated the anion secretory response of the endogenously expressed, native TMEM16A in CF-HBE to the P2Y2 receptor agonist, UTP (Figure 1). ETX001 treatment increased the sensitivity of the epithelium to UTP and enhanced the maximum efficacy (Figure 1A,B). The activity of ETX001 was independent of the duration of treatment, with both a 5 min (apical) and 96 h (basolateral) exposure potentiating TMEM16A function to a similar degree (Figure 1B). The amiloride-insensitive ISC was significantly enhanced following the 96 hour treatment with ETX001 (5.0 ± 0. 4 μA/cm^2^ in control versus 9.7 ± 0.4 μA/cm^2^ in the 96h ETX001 group; P=0.002, n=6 inserts per group) and was sensitive to inhibition by Ani9 (data not shown). These studies were performed in 3 donors of CF-HBE, including a code that was homozygous for ^F508del^CFTR.

**Figure 1.**
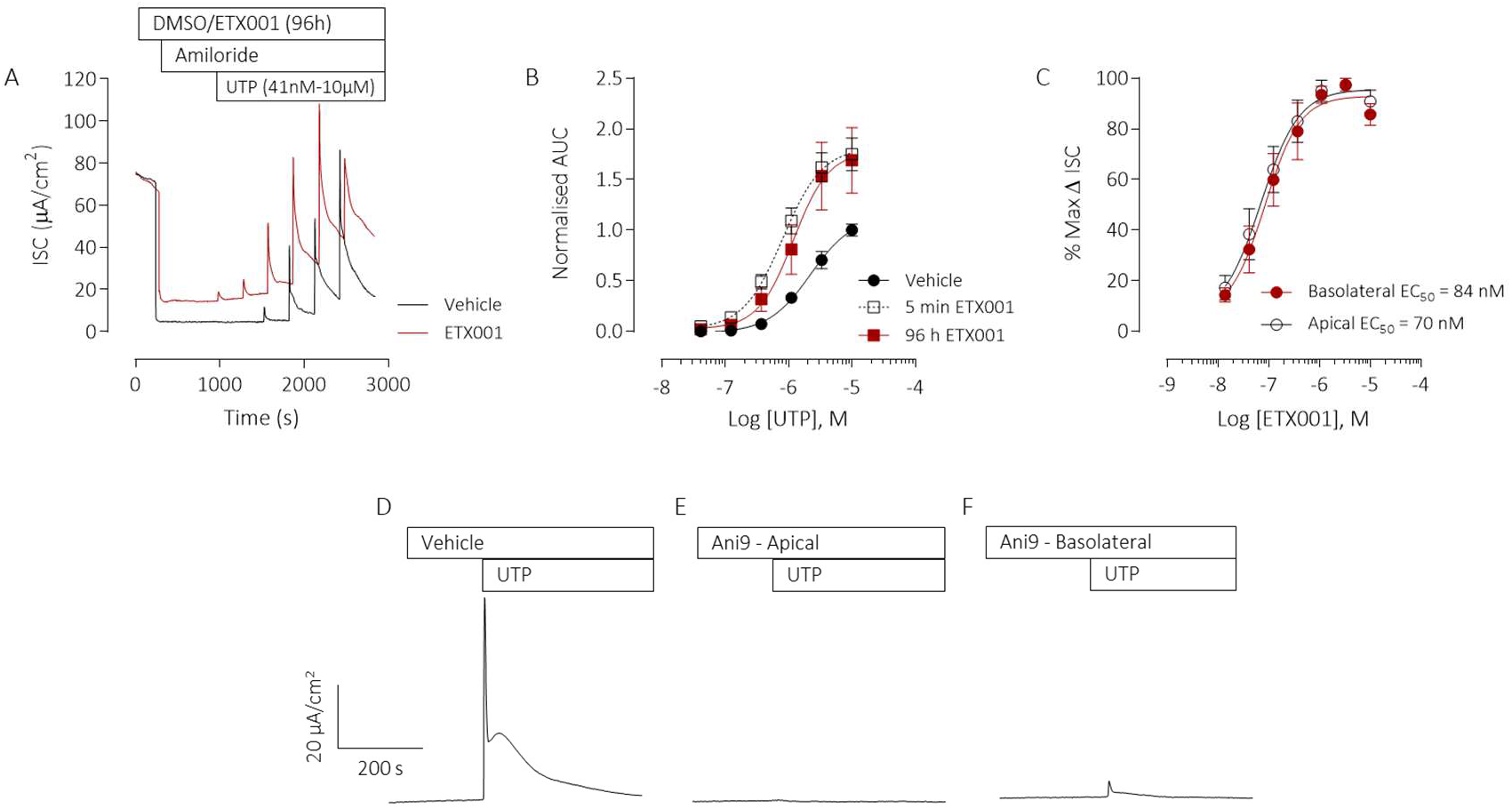
ETX001 potentiates the calcium activated anion secretory current in primary CF-HBE cells. Sample raw data trace illustrating the effects of a 96h incubation of CF-HBE with ETX001 (1μM; basolateral; red line) or vehicle (0.1% DMSO; grey line) on a UTP-stimulated anion current response (A). After inhibition of the ENaC current with amiloride (10μM) increasing concentrations of UTP were added to the apical side of the epithelium in a cumulative manner. Mean data (± SEM) illustrating the potentiation of the UTP concentration-response by ETX001 (96h or 5 min incubation) (B), normalised to the vehicle control (n=6-12 inserts per group). ETX001 potentiated the TMEM16A-mediated ISC in CF-HBE with a similar potency when added to either the apical or basolateral sides of the epithelium (C) (n=3-4 inserts per group). Sample raw data traces illustrating typical UTP-stimulated ISC responses of CF-HBE following pretreatment with vehicle (0.1% DMSO) (D), or with Ani9 (10 μM; apical) (E) or basolateral (F). All HBE had been pre-treated with IL-13 (10 ng/mL) for 48h prior to ion transport assay. CF-HBE from 3 independent donor codes were used for these studies (KK002C, KK036H, KK026K).

Following treatment of CF-HBE with CPA (2 μM), ETX001 induced a concentration-dependent increase in the short-circuit current. The potency and efficacy of ETX001 was independent of the sidedness of compound addition (Figure 1C). Similarly, the efficacy of the TMEM16A blocker Ani9, was independent of apical or basolateral addition. Figure 1D-F illustrates the attenuation of a UTP-stimulated anion-secretory response by Ani9 when administered to either the apical or basolateral side of the CF-HBE for 5 min before apical stimulation with the P2Y2-receptor agonist. These data validated the treatment of CF-HBE with either ETX001 or Ani9 by addition to the basolateral side of the epithelium in subsequent assays.

### Mucus secretion in vitro

Treatment of CF-HBE with ETX001 for 24 h failed to increase the secretion of either MUC5AC or MUC5B (Figure 2B,C). This was supported by TEM imaging and quantification of mucin granules in the goblet cells which likewise showed no evidence of an enhanced secretory response in the presence of the TMEM16A potentiator (Figure 2D,E). In contrast, treatment with PMA (300 nM) induced a significant increase in both the secretion of the mucins MUC5AC and MUC5B and also a reduction in the numbers of mucin granules in the secretory cells, consistent with the competence of the epithelium to undergo stimulated exocytosis of the goblet cells when suitably activated. In parallel experiments, cells were treated with IL-13 for 48h to upregulate TMEM16A expression and function as well as increasing MUC5AC protein expression (Figure 2A). IL-13 treatment significantly increased MUC5AC protein expression and reduced MUC5B expression but did not influence the lack of effect of ETX001 on mucin secretion or goblet cell exocytosis (Figure 2B,C). These studies were performed in 2 donors of CF-HBE, both homozygous for ^F508del^CFTR.

**Figure 2.**
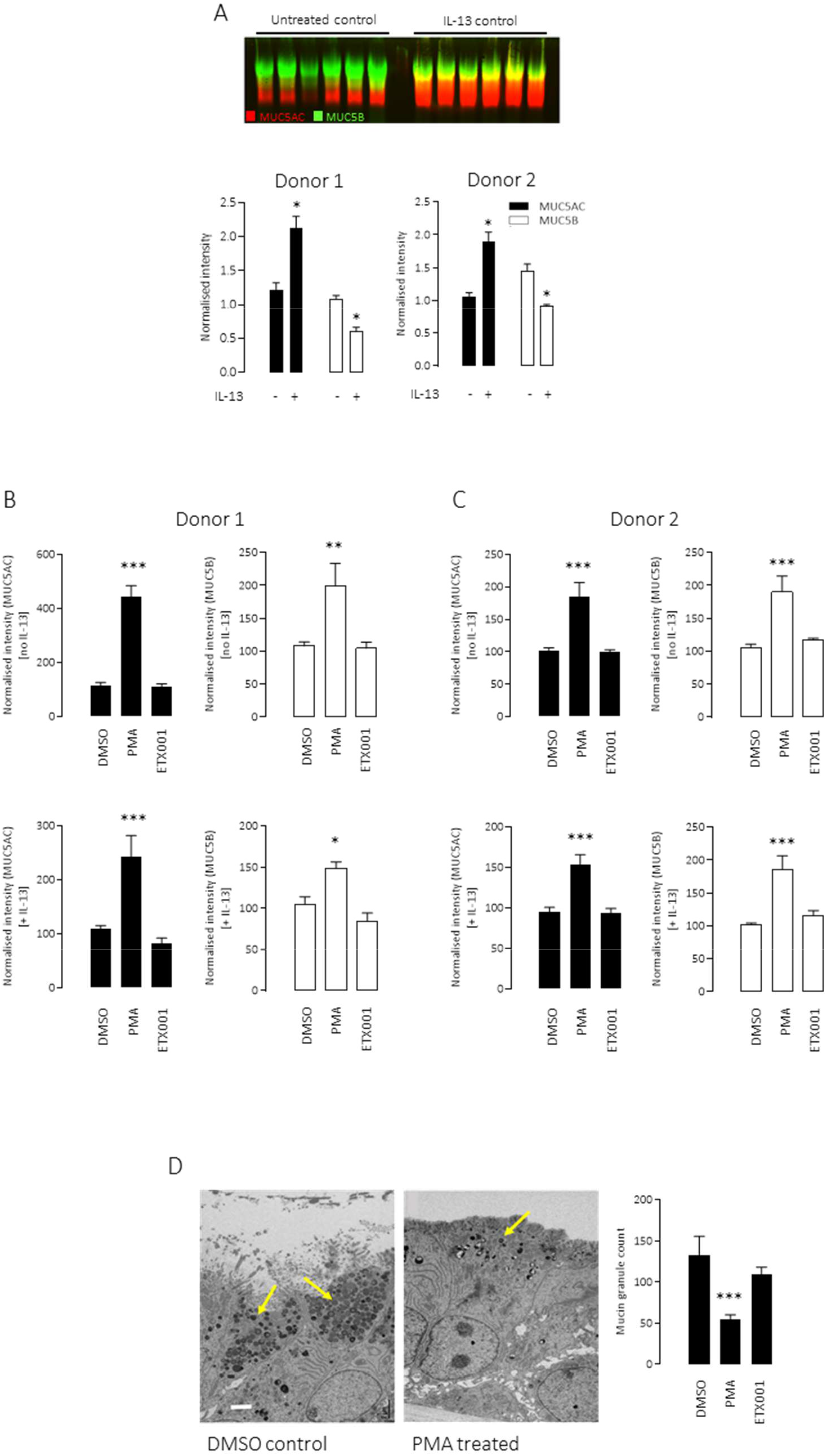
ETX001 does not enhance mucin secretion in primary CF-HBE cells. IL-13 treatment increases the mucosal accumulation of MUC5AC but reduces MUC5B accumulation in CF-HBE donor codes CFFT025K (Donor 1) and CFFT022L (Donor 2) (A). A sample Western blot of apical washings probed for MUC5AC and MUC5B is shown together with mean ± SEM data (n≥12 inserts per group). Panel (B) illustrates mean ± SEM data (n≥12 inserts per group; CFFT025K) for MUC5AC (filled bars) and MUC5B (open bars) accumulation recovered at 24h after treatment with either vehicle, PMA (300 nM) or ETX001 (1 μM). Top row are naïve inserts and lower row are IL-13 pre-treated. Panel (C) is the equivalent data-set obtained with donor code CFFT022L. Panel (D) illustrates sample TEM images of vehicle- and PMA-treated inserts combined with a graph showing the average mucin granule count per field of view. *P<0.05, ***P<0.0005 using one-way ANOVA with post-hoc Dunnett’s test.

### Mucus secretion in vivo

To avoid the potential of a bolus topical dose of a particulate suspension to influence goblet cell secretion, a metabolically stable analogue of ETX001, ETX004, was administered by i. p. injection to achieve sustained blood levels of a TMEM16A potentiator (Figure 3A). A solution formulation of ETX004 at doses of 0.3 and 3.0 mg/kg achieved blood exposures of approximately 3-25 fold over the EC_50_ for ETX004 on murine TMEM16A (Figure S1). ETX004, at these levels of exposure, did not stimulate any measurable emptying of goblet cell mucin content (Figure 3B,C) in contrast to the positive control ATP, a potent secretagogue, that induced a 45% reduction in goblet cell mucin content (P=0.001).

**Figure 3.**
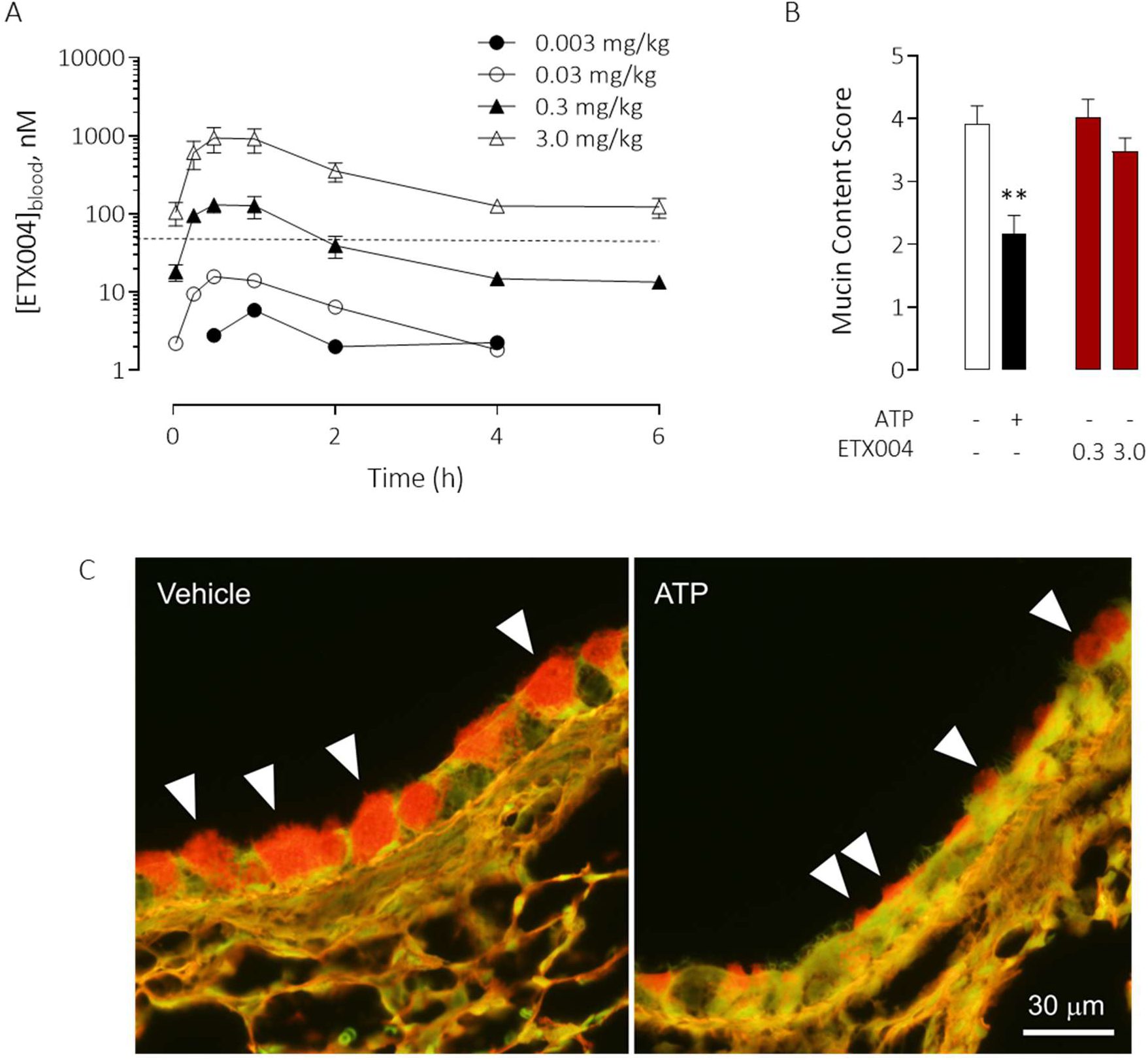
ETX004 does not stimulate airway goblet cell exocytosis in vivo. Following a single intraperitoneal dose, ETX004 was detected in blood in a dose-dependent manner to achieve concentrations in excess of the EC_50_ value for ETX004 on murine TMEM16A (38 nM; dotted line) at the 0.3 and 3.0 mg/kg doses (A). Mean data ± SEM are shown (n=3 mice per dose group). On a background of an IL-13 induced goblet cell metaplasia, ETX004 (0.3 and 3.0 mg/kg i.p.) was without effect on the emptying of airway goblet cells (B). The positive control, inhaled ATP, induced a significant emptying of goblet cells (B,C). A sample histology image illustrating the presence of PAFS stained material in airway goblet cells (white arrowheads) in vehicle treated mice compared with the reduced staining after aerosol exposure to ATP to induce exocytosis. Mean data ± SEM (n=10 mice per group). **P=0.001 using a Holm-Sidak’s multiple comparison test.

### Goblet cell formation in vitro

HBE cultured at air-liquid interface for 14-21 days (in Ultroser G-based ALI media) formed a well-differentiated epithelium, evidenced by the presence of both acetylated α-tubulin positive ciliated cells and MUC5AC positive goblet cells (Figure 4). Treatment of these cells with IL-13 for 96h significantly increased the MUC5AC positive stained area, consistent with an increase in goblet cell numbers. The parallel treatment of these cells with ETX001 failed to increase the MUC5AC positive stained area (Figure 4B-D). Similarly, co-treatment with both ETX001 and IL-13 failed to influence the MUC5AC positive stained area beyond the effect of IL-13 alone (Figure 4B-D). Treatment of cells with the TMEM16A blocker Ani9 was also without effect on the MUC5AC positive stained area in either the normal or IL-13 treated HBE (Figure 4B-D). The ciliated cell population was unaffected by any of the treatments based on the acetylated α-tubulin stained area (area data not shown).

**Figure 4.**
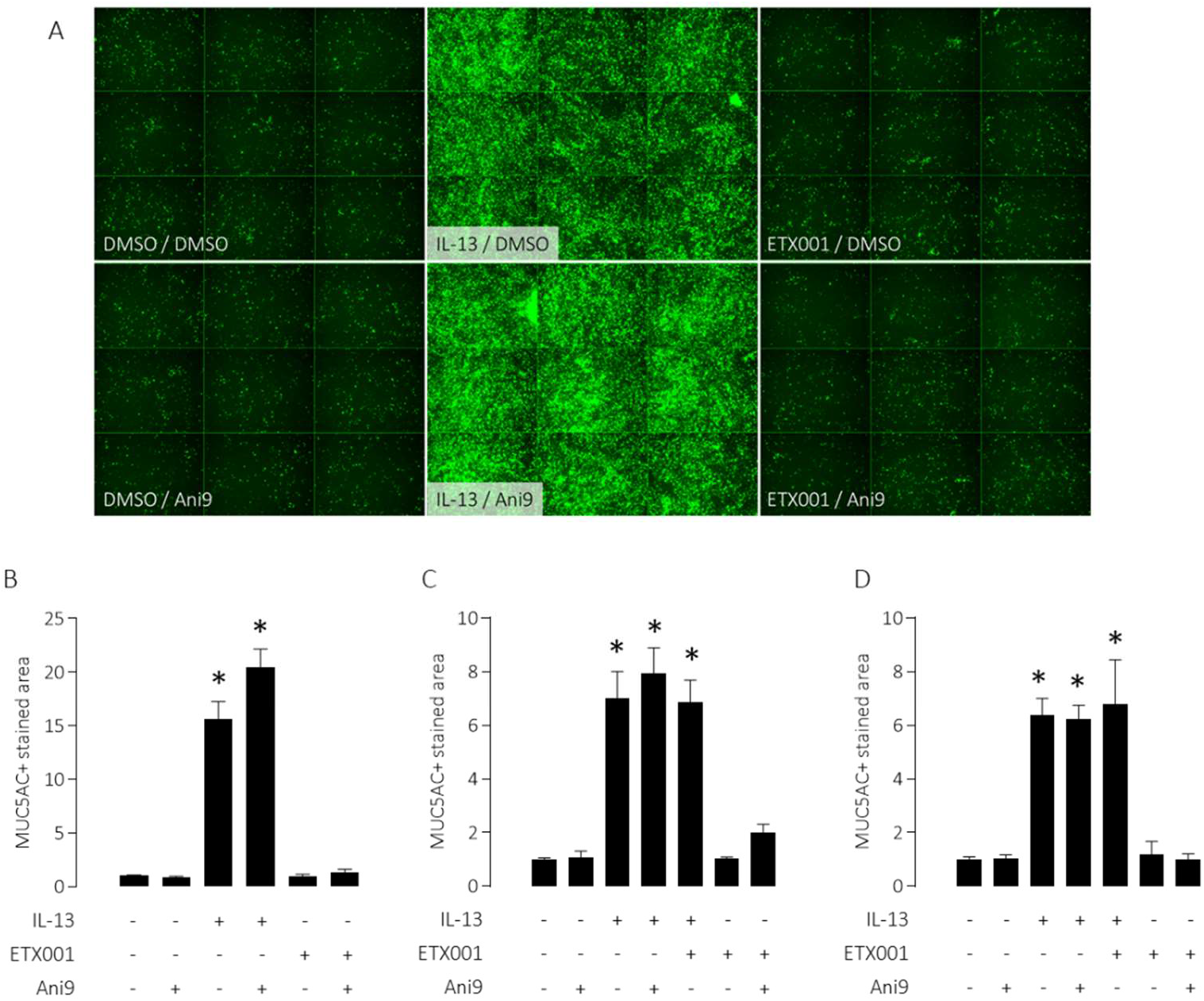
Neither potentiation nor inhibition of TMEM16A affects goblet cell numbers in primary HBE cells. TMEM16A modulators do not influence MUC5AC staining in HBE. MUC5AC positive goblet cells were quantified in 3 independent codes of non-CF HBE following treatments with IL-13, Ani9 (10,000 nM) or ETX001 (1 μM). Sample immune-stained images (green = MUC5AC+ cells) reflecting 9 images per individual inserts (A). Panels (B-D) illustrate mean MUC5AC+ stained area ± SEM (n≥4 inserts per group) normalized to the untreated control for each of the 3 donor codes. A one-way ANOVA with post-hoc Tukey’s test was used to test for significant differences between groups (*P<0.001 compared with the untreated control group).

### Smooth muscle contraction ex vivo

Freshly isolated human airways under an imposed tone showed a concentration-dependent relaxation to the positive control isoprenaline (Figure 5A). ETX001 failed to induce any change in airway tone, either relaxation or constriction. In contrast, the TMEM16A blocker niclosamide (10 μM) did significantly relax the tissues, although Ani9, a more potent and structurally diverse TMEM16A blocker did not have any effect. All airways studied had previously responded to carbachol challenge with a bronchoconstrictor response (data not shown).

**Figure 5.**
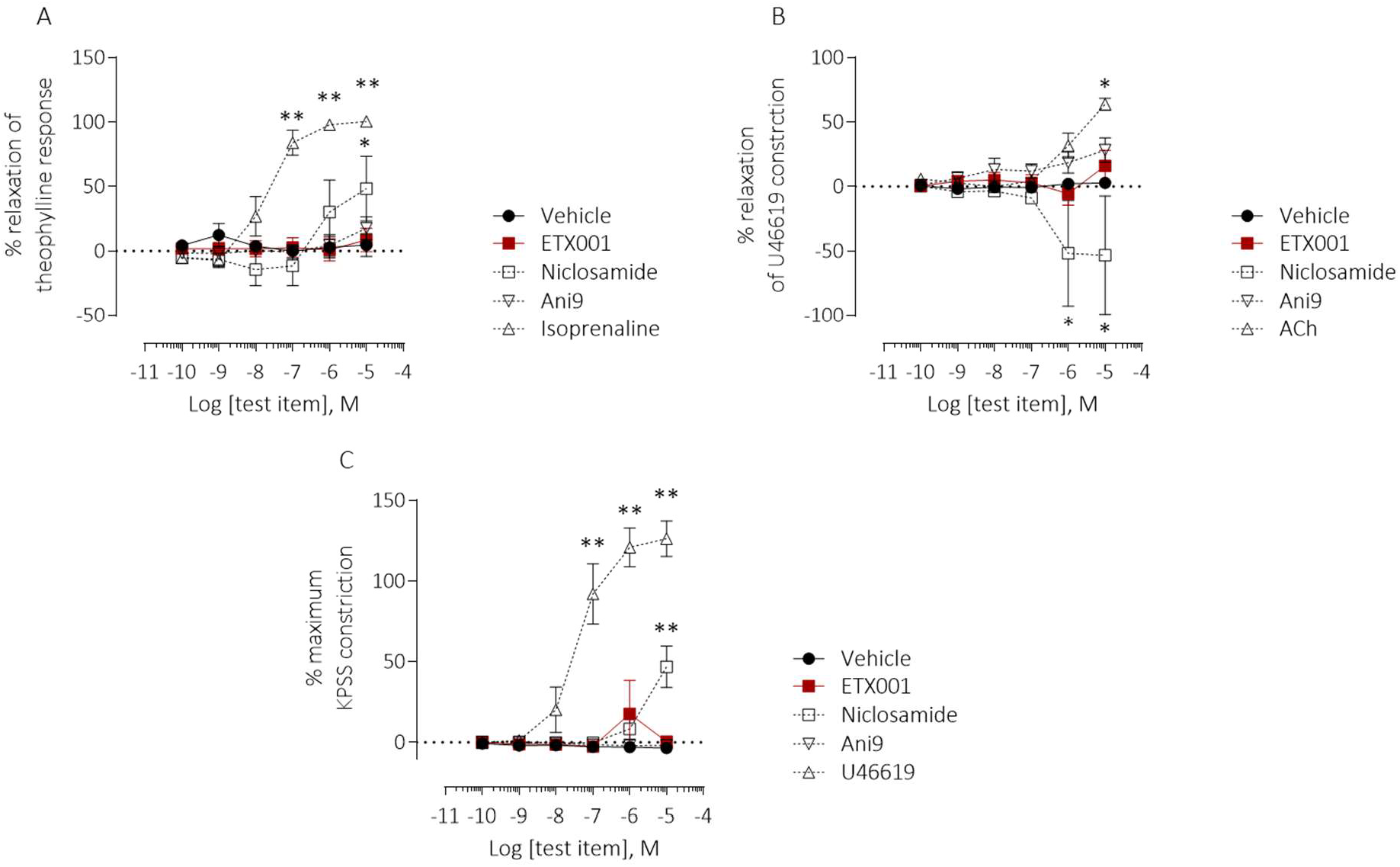
Neither potentiation nor inhibition of TMEM16A affects contractile or relaxant responses of human bronchi or human pulmonary preparations ex vivo. Effects of ETX001, Ani9 and niclosamide on isolated human airways (A) and pulmonary artery preparations in relaxant (B) or constrictor (C) format using wire myography. Mean data ± SEM are shown (n=3-5 replicates per group from individual donated lungs). A two-way ANOVA with post-hoc Dunnett’s test was used to test for differences between test items and vehicle (*P<0.01, **P<0.0001).

Freshly isolated human pulmonary arteries showed no response to ETX001 when studied in either the relaxation or constriction modes, although positive controls showed the predicted efficacy (Figure 5B,C). Niclosamide (10 μM) induced a vasoconstrictor response in both assay formats, an effect that was not evident with Ani9 treatment.

### Respiratory and cardiovascular function in vivo

ETX001 potentiated rat TMEM16A with an EC_50_ value of 96 nM (Figure S2). When dosed by intra-tracheal administration to telemetered rats, there were no ETX001-induced effects on either respiratory (Figure 6) or cardiovascular function (Figure 7). At the highest dose studied (2.5 mg/kg) there was a significant reduction in minute volume (Figure 6C), but only at a single, isolated timepoint (1.5h) with no evidence of time-dependent response as there were no trends at the timepoints either side (1.0 and 2.0h). Respiratory rate and tidal volume were not different from the values in the vehicle control group at any time during the study. There was a small (4%) decrease in diastolic blood pressure measured at a single timepoint (0.5h) in the 2.5 mg/kg treated group (Figure 7B). Again, there was no evidence of a time-dependent response.

**Figure 6.**
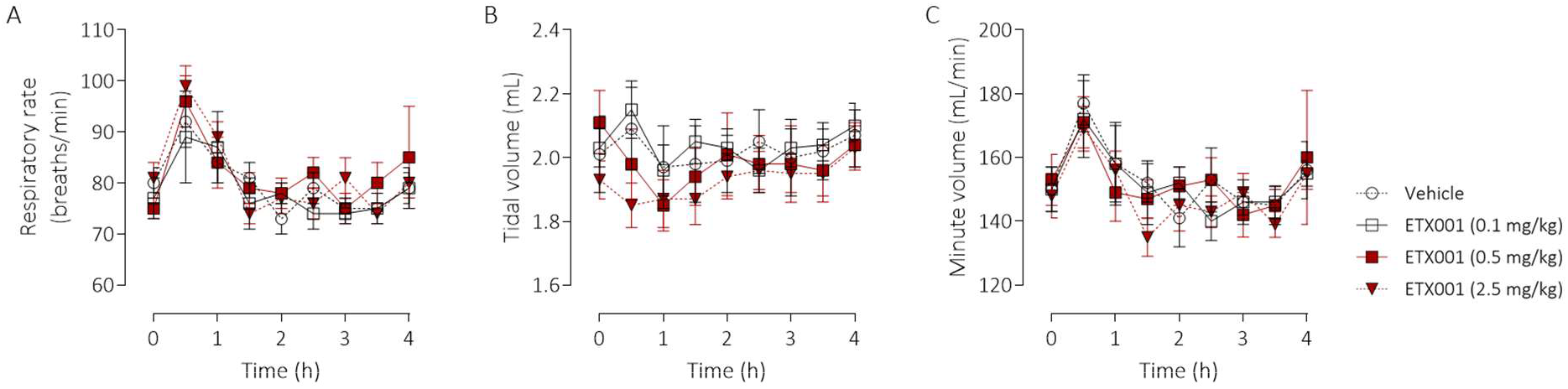
ETX001 has no effect on lung function in rat in vivo. The effects of ETX001 (0.1 – 2.5 mg/kg) or vehicle (0.05% Tween-80 in normal saline) on respiratory rate (A), tidal volume (B) and minute volume (C) when dosed by intra-tracheal instillation to male SD-rats. Mean data ± SEM (n=8 animals per group).

**Figure 7.**
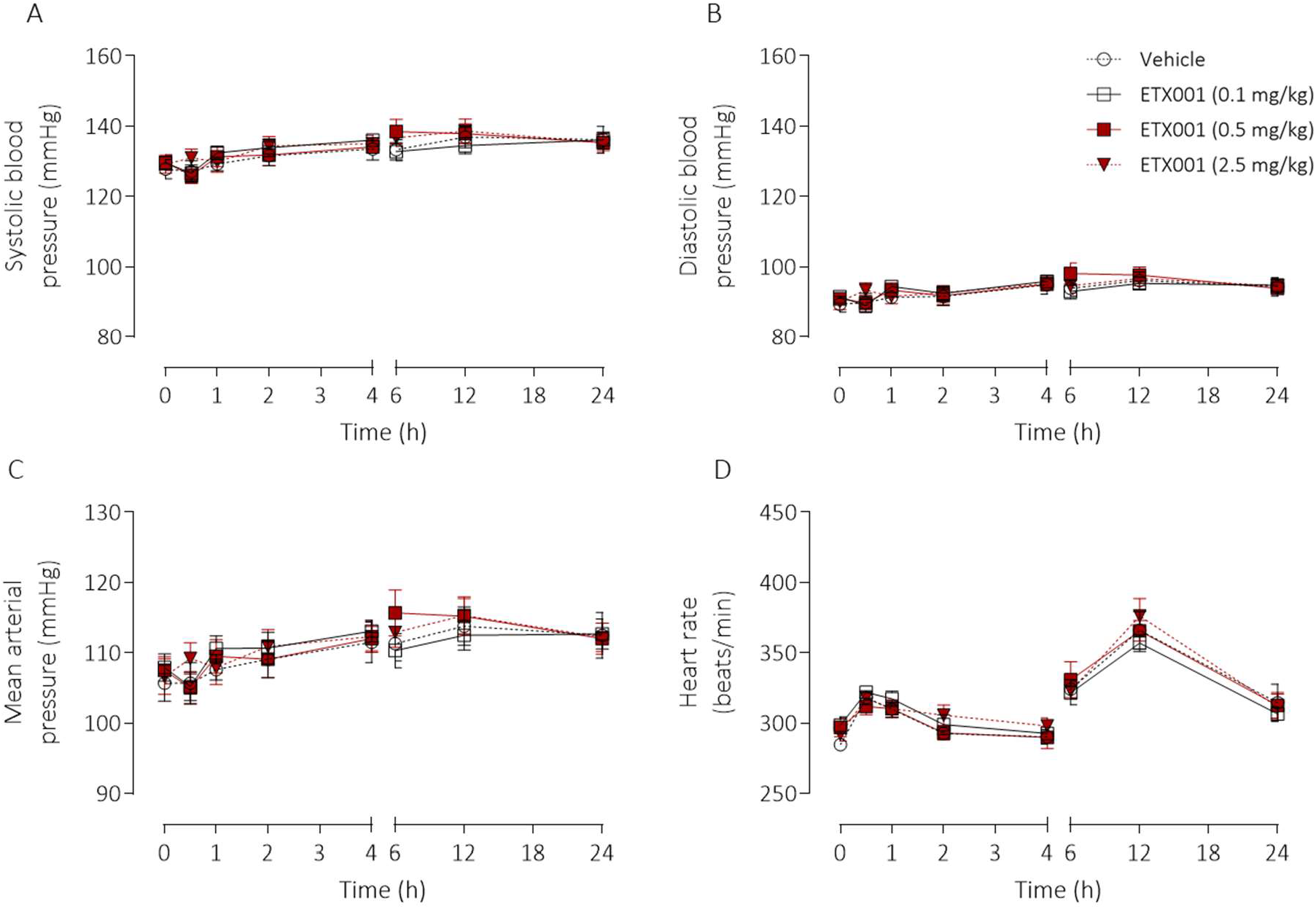
ETX001 has no effect on cardiovascular function in rat in vivo. The effects of ETX001 (0.1 – 2.5 mg/kg) or vehicle (0.05% Tween-80 in normal saline) on systolic (A), diastolic (B) and mean arterial (C) blood pressures together with associated changes in heart rate (D) when dosed by intra-tracheal instillation to male SD-rats. Mean data ± SEM (n=8 animals per group).

## Discussion

Since the identification of TMEM16A as the gene encoding the native CaCC, a number of physiological roles have been validated and expanded upon using a combination of genetic models and pharmacological modulators. Of relevance to the lung, TMEM16A loss-of-function studies have reported effects on goblet cell number and function (Benedetto et al., 2019; Huang et al., 2012; Zhang et al., 2015; Lin et al., 2015; Qin et al., 2016) as well as relaxation of airway smooth muscle (Danielsson et al., 2015, 2020; Huang et al., 2012; Miner et al., 2019) culminating in the recent proposal that inhibiting TMEM16A function would be of benefit in patients with respiratory diseases associated with mucus obstruction (Kunzelmann et al., 2019). Using the TMEM16A potentiator compounds ETX001 and ETX004, we have tested the hypothesis that positive modulation of the airway CaCC will induce goblet cell exocytosis, increase goblet cell numbers and cause airway and pulmonary arterial smooth muscle constriction. Despite illustrating the clear in vitro efficacy of ETX001 (Figure 1 for human data; Figure S2 for rat data) and ETX004 (Figure S2 for murine data) as potentiators of TMEM16A, we have not observed any effects on goblet cell number or function or on airway or pulmonary arterial smooth muscle contractility.

The functional upregulation of the airway CaCC by IL-13 in primary HBE, and under conditions that also induced a goblet cell metaplasia, was originally considered to be a coordinated and complimentary physiological response (Atherton et al., 2003; Scudieri et al., 2012). An increase in mucin production would require an increased fluid secretory capacity of the airway to maintain adequate hydration of the mucus gel. TMEM16A is abundantly expressed in goblet cells together with NKCC1 (Dolganov et al., 2001), the chloride transporter required to support vectorial chloride secretion across the airway epithelium. Having an apically located CaCC (TMEM16A) together with NKCC1 in the basolateral membrane supports a central role for a TMEM16A-dependent anion-secretory response mediated through the goblet cell. Recently however, additional roles for TMEM16A in goblet cell biology have been proposed.

A potential role of TMEM16A as being a key driver of the IL-13 induced goblet cell metaplasia has been suggested by several groups (Zhang et al., 2015; Lin et al., 2015; Qin et al., 2016). These studies have used non-selective blockers of TMEM16A such as niflumic acid, benzbromarone and T16Ainh-A01 to demonstrate a reduction in MUC5AC expressing goblet cell numbers in various airway epithelial cell models. However, the lack of effect of the potent TMEM16A blocker Ani9 in the present study is in direct contrast to these published studies and does not support a role for TMEM16A in the formation of goblet cells under IL-13 stimulation. The lack of efficacy of Ani9 on goblet cell formation cannot be due to a lack of access to target, as Figure 1D-E clearly demonstrates that Ani9 effectively blocks TMEM16A function in this cell system. The lack of effect of the TMEM16A potentiator ETX001 on goblet cell numbers further challenges the proposed role for TMEM16A channel function as a driver of differentiation towards the mucin producing phenotype. Consistent with our observations, a recent publication has also suggested that the upregulation of MUC5AC and TMEM16A are independent of one another and that there is no role for TMEM16A in goblet cell formation (Simões et al., 2019). One potential explanation for the disconnect between the Simões (2019) report and our present study with the literature may relate to the specificity of the TMEM16A blockers widely used in the earlier studies to establish the phenotype. For example, niflumic acid is a non-selective chloride channel blocker (Scott-Ward et al., 2004), inhibiting cyclo-oxygenase and several potassium channels (Busch et al., 1994; Hu et al., 2010; Walker et al., 2002). T16Ainh-A01 and benzbromarone likewise have additional pharmacological (Boedtkjer et al., 2015; Enomoto et al., 2002; Wang et al., 2007) and toxicological activities (Felser et al., 2014), that may have influenced the outcome of the published studies.

Attenuated goblet cell-derived mucin secretion secondary to the loss of expression of *Tmem16a* in mice has been described by Benedetto and colleagues (2019). When *Tmem16a* gene expression was specifically silenced in *Foxj1*^+^ multiciliated cells, the authors described an attenuated paracrine signalling mechanism to the goblet cell which inhibited exocytosis. Using well-differentiated HBE cultures in the present study, both multiciliated and goblet cells are represented and therefore presumably any putative paracrine signalling pathway. In this system, potentiation of TMEM16A with ETX001 failed to influence the secretion of mucus irrespective of pre-treatment with IL-13 to both increase the expression of TMEM16A and MUC5AC. The reported TMEM16A activator E_act_ has also been demonstrated to induce airway goblet cell emptying of stored mucus (Benedetto et al., 2019). Although originally reported to not directly influence [Ca^2+^]_i_, (Namkung et al., 2011) E_act_ has recently been demonstrated to induce a non-specific elevation of [Ca^2+^]_i_ which then drives an indirect activation of TMEM16A (Genovese et al., 2019). It therefore seems likely that any E_act_ induced emptying of airway goblet cells is due to a TMEM16A-independent elevation of [Ca^2+^]_i_, a well-documented mechanism for exocytosis (Lethem et al., 1993). Importantly, ETX001 was demonstrated to have no direct effect on [Ca^2+^]_i_, and also did not influence a P2Y2 receptor driven elevation of [Ca^2+^]_i_ (Danahay et al., 2020). These data would appear to challenge the direct linkage between the positive regulation of TMEM16A function and mucus release in human airway epithelial cells. In vivo, extended systemic exposure to a potent TMEM16A potentiator ETX004 also failed to demonstrate any evidence of goblet cell emptying. ETX004 (i.p.) achieved sustained blood levels that were in excess (up to 25x) of the on target EC_50_. These data further support the lack of a role for TMEM16A in the regulation of goblet cell exocytosis.

The role of the CaCC in smooth muscle contractility has been well documented to contribute towards depolarisation of the plasma membrane and subsequent Ca^2+^ influx. Huang et al., 2012 implicated a role for TMEM16A in the regulation of murine airway smooth muscle function and more recently it has been demonstrated that non-selective TMEM16A blockers including benzbromarone, T16Ainh-A01 and niclosamide relax airway smooth muscle preparations (Danielsson et al., 2015, 2020; Miner et al., 2019). The reported TMEM16A activator E_act_, has also been shown to induce airway smooth muscle contraction and bronchospasm in vitro and in vivo (Danielsson et al., 2015, 2020). The present study has confirmed the broncho-relaxant effect of niclosamide in isolated human bronchi. However, the lack of effect of both Ani9 and ETX001 in this system challenges a role for TMEM16A in the regulation of human airway smooth muscle contractility. Furthermore, when dosed directly into the airways of rats, ETX001 failed to show any robust effects on lung function parameters. The doses of ETX001 used in this study (0.5 – 2.5 mg/kg) were in excess of the doses administered to conscious sheep (35 – 100 μg/kg) to observe efficacy in a model of mucociliary clearance (Danahay et al., 2020). Given that ETX001 shows good species cross-reactivity in rat and was dosed at >70x the efficacious, durable dose in the sheep, it seems unlikely that the lack of effect of ETX001 would be due to inadequate exposure of the airway smooth muscle to compound. A potential explanation for the inconsistency between the present data with ETX001 and Ani9 and published studies may again relate to the lack of selectivity of the TMEM16A blockers and activator, E_act_, that have been widely used in the literature. Of particular note, niclosamide is a mitochondrial toxin within the same concentration range required to block TMEM16A (Tao et al., 2014). As discussed above, E_act_, is not a direct TMEM16A activator but rather induces an elevation of [Ca^2+^]_i_ that would induce smooth muscle contraction, irrespective of any putative effect on TMEM16A.

Roles for TMEM16A in the regulation of vascular smooth muscle have been similarly proposed although the function is controversial (Leblanc et al., 2015). To this end, we have been unable to observe any effects of either Ani9 or ETX001 in isolated human pulmonary artery preparations. Niclosamide induced a contractile effect in both assay formats evaluated, although a role for TMEM16A inhibition cannot be ascribed for the reasons relating to selectivity outlined above. ETX001 also failed to influence cardiovascular function in vivo following local delivery into the airways at doses up to and including the highest dose studied (2.5 mg/kg). However, it should be noted that ETX001 has been designed as an inhaled drug with high plasma clearance and short plasma half-life that will limit systemic exposure.

The data presented in the current study are therefore inconsistent with the hypothesis that the positive modulation of TMEM16A will provoke goblet cell exocytosis, goblet cell metaplasia, induce bronchoconstriction or cause pulmonary artery constriction. Furthermore, inhibition of TMEM16A with the potent and selective channel blocker Ani9, did not relax human bronchial smooth muscle, influence goblet cell function or alter pulmonary artery tone, phenotypes previously observed with non-selective blockers. Together, these data challenge the potential therapeutic utility of TMEM16A blockers for the treatment of respiratory diseases (Kunzelmann et al., 2019) and support the evaluation of TMEM16A potentiators as an approach to hydrate mucus in diseases such as cystic fibrosis (Kunzelmann et al., 2012; Mall & Galietta, 2015; Sondo et al., 2014).

