## Supplemental information for "Potentiating TMEM16A channel function has no effect on airway goblet cells or bronchial and pulmonary vascular smooth muscle function"

### Methods

#### Cross-species TMEM16A potentiator pharmacology

Cell lines stably expressing full-length variants of either mouse (NM_178642.5) or rat (NM_01107564.1) were used (Scottish Biomedical, UK). Cells were cultured in Ham's F12 medium supplemented with 10% foetal bovine serum, 2mM L-glutamine, 100 U/ml penicillin, 100 µg/mL streptomycin and 250 µg/mL hygromycin B. Cells were maintained at 37°C in 5% $CO_2$ in air incubator and grown to 70-80% confluence prior to passaging or use in electrophysiology experiments. Whole-cell patch clamp recordings were made as previously described (Danahay et al.,2020) using a QPatch automated electrophysiology platform. In brief, currents were quantified using a combined step (-70 mV to +70 mV) and voltage ramp (-90 mV to +90 mV) protocol in experiments using anion selective recording solutions to isolate the TMEM16A current. The extracellular recording solution consisted of (mM) 130 NMDG-Cl, 2 $CaCl_2$, 1 $MgCl_2$, 10 HEPES with the pH adjusted to 7.3 with HCl and the osmolarity to 320-330 mOsm with sucrose. The intracellular recording solution consisted of (mM) 130 NMDG-Cl, 1 $MgCl_2$, HEPES 10, EGTA 20, BAPTA 10, Mg-ATP 2 with $CaCl_2$ added to provide a free $[Ca^{2+}]_i$ experimentally derived to be the $EC_{20}$ for TMEM16A channel activation. For the CHO-mTMEM16A (murine) rTMEM16A (rat) this was 260 nM and 415 nM respectively. The effect of test compounds on TMEM16A currents was quantified as the percentage change in the basal current level (activation) at the end of the +70 mV step. The compound effect was analysed by calculating the percentage change in current relative to the starting baseline current. The compound effect was always measured as the maximum current change within each concentration incubation. Concentration-response curves were generated using these values and fitted using a Hill plot within the QPatch software to determine the $EC_{50}$ value.

#### In vivo pharmacokinetic profile of ETX004

Male C57bl6 mice (26-32 g) were dosed with ETX004 (by intraperitoneal injection. ETX004 was dissolved in 5% NMP/95% HPCD (20%) and dosed at 0.003 to 3.0 mg/kg (10 mL/kg). At regular intervals after dosing, blood was sampled from the tail vein (25 µL) and diluted 1:1 with water. Diluted blood was stored at -20°C prior to bioanalysis. ETX004 concentrations in blood were determined using standard LCMS-MS methods.

### Results

#### Cross-species TMEM16A potentiator pharmacology

ETX004 potentiated murine TMEM16A activity to a maximum 210% efficacy and with an $EC_{50}$ value of 38 nM (n=6) (Figure S1). ETX001 potentiated rat TMEM16A activity to a maximum 337% efficacy and with an $EC_{50}$ value of 88 nM (n=41) (Figure S2).

#### In vivo pharmacokinetic profile of ETX004

ETX004 (0.3 and 3.0 mg/kg i.p.) achieved sustained blood levels reaching $C_{max}$ values of 131 and 939 nM respectively, with $T_{max}$ of 30 min.

### Conclusions

ETX004 and ETX001 show a potent potentiation of murine and rat TMEM16A respectively. Furthermore, potency and efficacy of ETX001 are similar to the values previously described for this compound on the human channel (Danahay et al., 2020).

ETX004 can be dosed by i.p. injection to mice at dose levels that achieve systemic exposures of parent compound up to 25-fold higher than the $EC_{50}$ for murine TMEM16A.

Supplementary figure legends

Figure S1 ETX004 potentiates murine TMEM16A in vitro

Potentiation of recombinant murine TMEM16A currents by ETX004 as assessed by automated patch electrophysiology. Sample data trace showing potentiation of mTMEM16A current by ETX004 at the indicated concentration. Voltage protocol used to generate the recording is shown above; dotted line indicates the zero-current level. Concentration-response curve for ETX004 (n=6). Symbols are the mean ± sem.

Figure S2 ETX001 potentiates rat TMEM16A in vitro

Potentiation of recombinant rat TMEM16A currents by ETX001 as assessed by automated patch electrophysiology. Sample data trace showing potentiation of rTMEM16A current by ETX001 at the indicated concentration. Voltage protocol used to generate the recording is shown above; dotted line indicates the zero-current level. Concentration-response curve for ETX001 (n=41). Symbols are the mean ± sem.

Supplementary figures

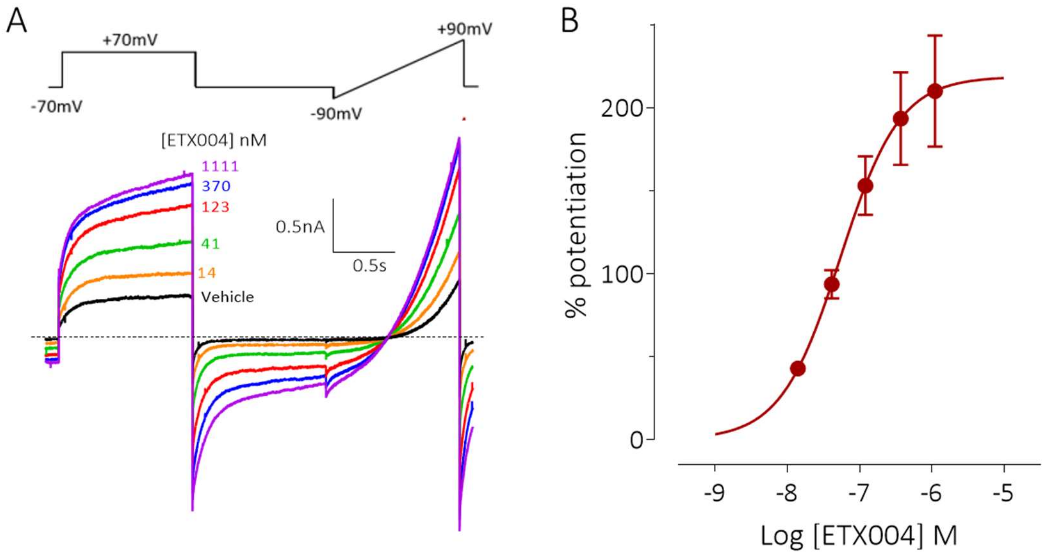


Figure S1

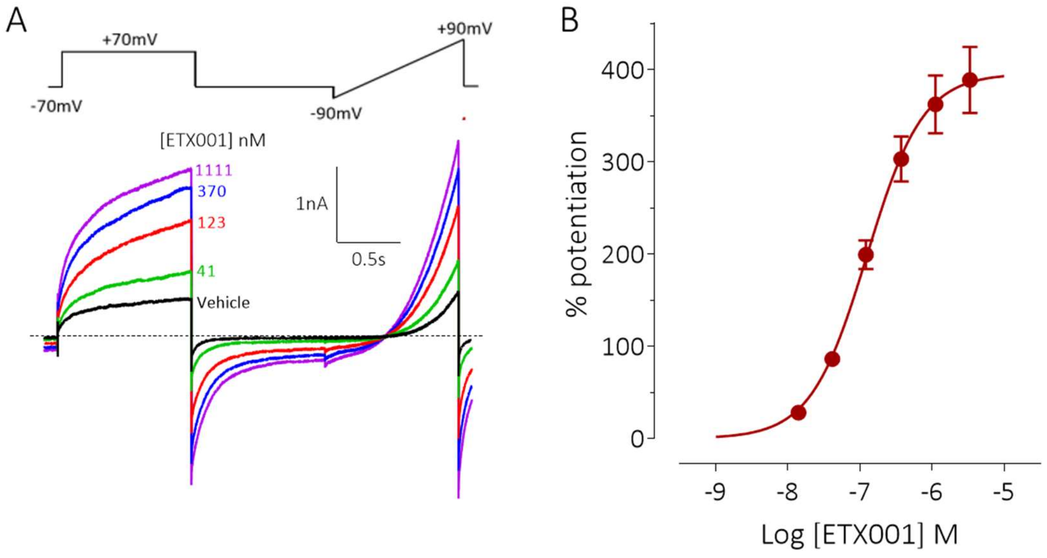
A
+70mV
+90mV
-70mV
-90mV
[ETX001] nM
1111
370
123
41
Vehicle
1nA
0.5s
B
% potentiation
400
300
200
100
0
-9
-8
-7
-6
-5
Log [ETX001] M

Figure S2
